# SAMCell: Generalized Label-Free Biological Cell Segmentation with Segment Anything

**DOI:** 10.1101/2025.02.06.636835

**Authors:** Alexandra D. VandeLoo, Nathan J. Malta, Emilio Aponte, Caitlin van Zyl, Danfei Xu, Craig R. Forest

## Abstract

**Background:** When analyzing cells in culture, assessing cell morphology (shape), confluency (density), and growth patterns are necessary for understanding cell health. These parameters are generally obtained by a skilled biologist inspecting light microscope images, but this can become very laborious for high throughput applications. One way to speed up this process is by automating cell segmentation. Cell segmentation is the task of drawing a separate boundary around each individual cell in a microscope image. This task is made difficult by vague cell boundaries and the transparent nature of cells. Many techniques for automatic cell segmentation exist, but these methods often require annotated datasets, model retraining, and associated technical expertise.

**Results:** We present SAMCell, a modified version of Meta’s Segment Anything Model (SAM) trained on an existing large-scale dataset of microscopy images containing varying cell types and confluency. We find that our approach works on a wide range of microscopy images, including cell types not seen in training and on images taken by a different microscope. We also present a user-friendly UI that reduces the technical expertise needed to use this automated microscopy technique.

**Conclusions:** Using SAMCell, biologists can quickly and automatically obtain cell segmentation results of higher quality than previous methods. Further, these results can be obtained through our custom GUI without expertise in Machine Learning, thus decreasing the human labor required in cell culturing.

## 1 Background

Cell culture is widely used for biological research applications. In conducting cell culture, microscopy is used to assess cell attributes like confluency (density of cell growth), morphology (cell shape), and count - metrics which relate to cell health and allow a researcher to assay viability visually throughout the cells’ growth period ^1^. These parameters, typically obtained manually by a skilled biologist, are important to the success of biological experiments. Confluency, for example, is often used to gauge when cells are ready to be passaged, differentiated, or otherwise manipulated ^2^ and requires experience to assess accurately by eye.

Automating this process is an open area of research. Automatically segmenting cells is a difficult problem. Data annotation is especially laborious as cells routinely number in the hundreds per image. Further, due to differences in imaging parameters like contrast and field brightness, high variance in pixel intensity indicative of cell boundaries exists between images from different microscopes even of the same imaging technique (e.g., Phase-Contrast, Brightfield). Finally, cells are often tightly clumped together and have difficult to discern edges even for annotators, as illustrated in Figure 1. An ideal model would accurately predict cell boundaries for a wide range of images and cell types to reduce the need for manual data annotation. If accurate cell boundaries could be produced, relevant parameters to cell health like average number of nearest cell neighbors, average cell aspect ratio, and average cell size could be automatically calculated and reported. Reliably automating the segmentation of microscopy images across cell types and imaging parameters has not yet been achieved ^3^.

**Fig. 1:**
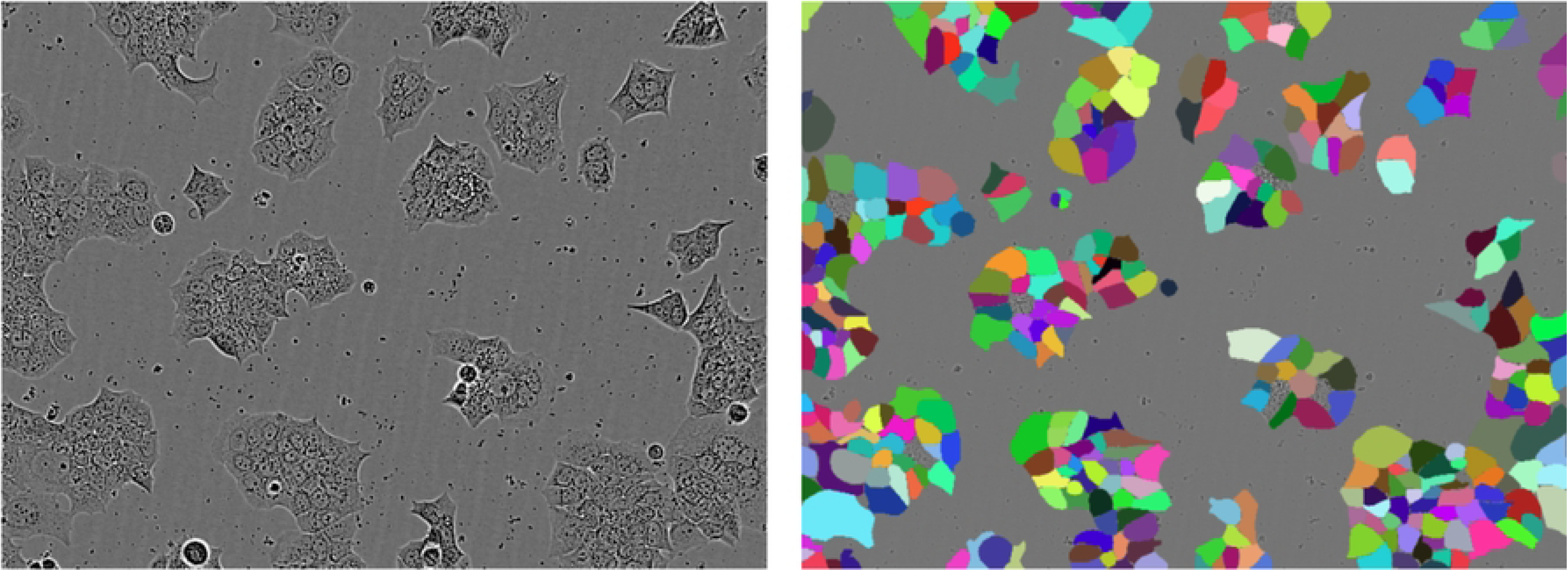
(a) An image from the LIVECell Test Set containing clumped cells with hard to distinguish edges. Note: image contrast is artificially enhanced for the reader. (b) overlaid ground-truth masks

Driven by the needs of cell culturing, we are concerned with finding the boundaries of entire, unlabeled cells in this work. Unlike other approaches ^4^, we do not explore segmenting the nucleus or other regions as separate from the remainder of the cell.

Interestingly, cell segmentation poses new challenges not seen in standard, multi-class segmentation like the previous Segment Anything Model (SAM) work ^5^. For example, cells are often packed tightly in microscope images, with weak or even ambiguous edge features. As such, care must be taken not to merge adjacent cells in predicted masks. One approach, employed by the introductory U-Net paper, is to segment the image into two categories, “cell” and “background”, with a minimum 1-pixel border of category background between adjacent cells. Then, connected components of category “cell” are extracted from the prediction as individual cells. To avoid clumped cells being merged by the model, a pixel-wise weighted loss function with a higher weight near cell boundaries is used ^6^. Unfortunately, due to the constraint that predicted cell masks cannot be in contact with each other, this approach restricts the model’s ability to accurately predict cell mask when dealing with clumped cell masks. Further, the light-weight U-Net model can under-fit larger datasets and has a reduced potential to benefit from pretraining compared to larger models.

Subsequent work expands on this idea, introducing a 3rd “cell border” category and experimenting with various loss function weightings. After classification, these edge pixels are assigned to the nearest cell ^7^. Although this additional category eliminates the need for a background border, “cell border” category pixels can become difficult to assign to the correct cell when cells are not circular or contain fine features, like long offshoots. This approach still has limited benefit from pretraining and like the base U-Net has the potential to under-fit large datasets. From initial experiments with Segment Anything, we observe that for cells with weak edge features, a three-category classification with a weighted loss function is insufficient to prevent the merging of clustered cells.

Another prominent cell segmentation work, Cellpose, uses a modified U-Net architecture to estimate a specialized vector field, computed from a simulated diffusion process^8^. The authors define a diffusion process that produces a vector field where, for each cell, gradient vectors point away from the cell’s center in all directions. Cell masks are then computed by finding the fixed points of this vector field, locations where the vectors from neighboring cells intersect. Because Cellpose calculates and predicts this vector field independently for the x and y components, rotation in data augmentation becomes non-trivial. Finally, being based on U-Net, the model may under-fit larger datasets and has a reduced advantage from pretraining.

In April 2023, Meta released Segment Anything Model (SAM) ^5^, a general model for image segmentation. SAM was trained on a diverse dataset of 11 million everyday images containing, in total, more than 1 billion segmentation masks. Due to this extensive dataset, Segment Anything is a great generalist: showing strong performance even on datasets not seen during model training. Given a point or bounding box in an image (the “prompt”), SAM can produce a reasonable segmentation boundary. Alternatively, SAM can generate masks automatically, by uniformly sampling points around the image, and keeping the most confident segmentations. Thanks to the extensive dataset used, SAM can produce a reasonable boundary for a wide set of objects.

Because of SAM’s strong performance on natural images, there have been attempts to fine-tune it for the biomedical domain. MedSAM ^9^, for example, fine-tunes SAM using more than a million images collected from various medical imaging modalities, like X-Rays, Computed Tomography (CT) Scans, and ultrasounds. These authors preserve SAM’s prompting capability, training MedSAM to determine masks from a bounding box. In training, the authors completely update the weights in the image encoder and mask decoder, while freezing the prompt encoder. MedSAM delivers strong results, exceeding baselines in a variety of segmentation tasks and offering impressive zero-shot performance on unseen datasets. Preserving the prompt does allow a user to have greater control over the model’s segmentation. This prompt based approach is less useful for cell segmentation, however, because of the sheer number of cells in microscopy images. Drawing a bounding box or selecting a prompt point for a large number of cells would become incredibly laborious and limit the model’s usefulness to biologists.

SAMed ^10^, a concurrent work to MedSAM, fine-tunes SAM to segment individual organs in CT scans. Interestingly, SAMed does not require a prompt and, unlike SAM, can segment images into distinct categories, like liver, stomach and pancreas. Further, SAMed is trained on the comparably smaller scale Synapse multi-organ CT dataset, which consists of just a few thousand images. SAMed uses Low Rank Adaptation ^11^ to fine-tune SAM’s image encoder, while retraining all parameters in the mask decoder. SAMed’s ability to segment images without a prompt is a desirable characteristic for cell segmentation. However, because SAMed categorizes images into a discrete number of categories, we find that it still is prone to combining adjacent cells, like the U-Net approaches ^6, 7^.

Due to the popularity of Segment Anything Model, a few concurrent works also use it to approach the cell segmentation task. CellSAM ^12^, for example, first creates rectangular bounding boxes around each cell in an image with an object detection model. These bounding boxes serving as prompts for a fine-tuned SAM. Thus, CellSAM can produce cell boundaries while preserving the prompting capability of Segment Anything Model. Another work, “Segment Anything for Microscopy” ^13^ introduces an extension of the default Segment Anything Model, called micro sam. Like CellSAM, micro sam offers a version of Segment Anything Model fine-tuned on a dataset of microscopy tasks. Uniquely, however, micro sam offers support for some microscopy specific tasks: tracking cells over time in images and segmenting a cell in 3D collection of images. It also offers integration with an existing GUI for easy manual prompting by a user. Taking a different approach, CellViT ^14^ leverages the pretrained image encoder from Segment Anything, while replacing SAM’s mask decoder with a U-Net inspired segmentation decoder. CellViT is able to segment and classify cell nuclei in tissue images with impressive results.

Hoping to bring Segment Anything’s generalization to the field of cell segmentation, we were also motivated to adapt SAM to the new domain. We present SAMCell, a fine-tuned approach based on SAM for predicting cell boundaries in microscope images. Unlike the default SAM, we observe success in cases even when cells are densely packed or boundaries are soft. Like concurrent works, SAMCell inherits Segment Anything’s main advantages: a high parameter, Vision Transformer ^15^ based architecture and extensive pretraining. These advantages improve generalization and allow our method to better fit a large dataset, compared to other approaches. We take a unique approach to finetuning compared to other concurrent works. Rather than using an object detection model for prompting (like CellSAM) or a custom decoder module (like CellViT) we disregard prompting and pose segmentation as a regression task. We fine-tune SAM to output a real-valued distance map describing the euclidean distance to a cell border for each pixel in the image. We then recover the boundary using a post-processing technique based on the watershed algorithm.

We find that our method exceeds the performance of existing approaches on both cells similar to those seen in training (test-set) and on completely novel cell lines from images taken by other microscopes (zero-shot).

## 2 Methods

### 2.1 Datasets

We evaluate SAMCell for two main use cases. First, the case where a sufficiently large dataset is available to train SAMCell. After training, we can observe performance on the test set, a portion of the dataset not seen during training. In this case, the training images and test images for evaluation are similar. We also evaluate the “zero-shot” case, where no annotated dataset is available. In this other case, we train the model on a large dataset of cell images, with the aim of generalization. Then, we evaluate on images dissimilar to what the model has seen during training. We select datasets to evaluate both these “test set” and “zero shot” use cases.

#### 2.1.1 Large-Scale Datasets for Test-Set Evaluation

For model training and test set evaluation, we select two existing large-scale datasets of cell images. First, we choose the LIVECell dataset ^16^ which consists of more than 5000 phase-contrast images, across 8 cell types. The dataset consists of cells with a range of confluency and morphology. Further, many images have low contrast. Together, these two characteristics produce a difficult segmentation task which is useful in evaluating model performance. Unfortunately, all images in this dataset were produced with the same phase-contrast microscope. Even among phase-contrast microscopy, different microscopes can produce dissimilar pictures because of variations in imaging parameters like contrast, brightness, and resolution. As such, we conjecture that only including images from a single microscope limits LIVECell’s coverage of the cell segmentation task.

To complement LIVECell, we also employ the Cytoplasm dataset gathered by the authors of the Cellpose model ^8^. This dataset consists of microscopy images scraped from the internet which were subsequently annotated. Because of the nature of web scraping, these images came from a variety of different individual microscopes. Further, a variety of microscopy techniques are included in this dataset, including bright-field images, membrane-labeled cells, and fluorescent labeled proteins. This dataset contains around 600 images – a smaller-scale dataset than LIVECell. Despite this, we expect the Cytoplasm dataset to be better suited for training generalist models because of the wider range of images included. Some of the images in the Cytoplasm dataset are 3-channel, with separate colors showing the nucleus and membrane using fluorescent labeling. Because we are specifically concerned with label-free, whole-cell segmentation in this work, we convert all images in this dataset to grayscale for training and evaluation. Because images in the Cytoplasm dataset are of different sizes, we also resize all images to 512x512, preserving the aspect ratio by adding a border as needed. A majority of images are near this size and close to square. This resizing allows us to more easily train our model and baselines by avoiding jagged arrays.

#### 2.1.2 Small-Scale Datasets for Zero-Shot Evaluation

We produce two novel, small-scale datasets. Each contains 5 images of varied confluency and microscope contrast annotated by an expert biologist co-author. These datasets contain images of the Human Embryonic Kidney (HEK) 293 and Neuro2a (N2a) cell lines respectively. We name these datasets after our lab, the Precision Biosystems Laboratory (PBL), and the cell lines they contain, yielding *PBL-HEK* and *PBL-N2a* respectively.

Although these datasets are of small scale, they are sufficient for zero-shot evaluation as each image contains approximately 300 cells. These datasets are entirely used for testing, to evaluate how SAMCell and our baselines generalize to unseen images from different Phase-Contrast microscopes. We evaluate zero-shot performance using SAMCell and baseline models trained on the Cellpose Cytoplasm dataset, because of the higher diversity in microscopy techniques and cells imaged.

### 2.2 Evaluation Metrics

For evaluation, we employ two cell segmentation metrics used within the Cell Tracking Challenge ^17^, namely, Detection Accuracy Measure (DET) and Segmentation Accuracy Measure (SEG) ^18^. These metrics assess the models’ accuracy for Detection (the ability to identify objects correctly) and Segmentation (the ability to match the boundaries of each object correctly) respectively in comparison to a given “ground-truth” annotation. The SEG metric uses a simple Jaccard index: the intersection between a ground truth annotation and a corresponding model prediction, divided by the union of these two values. The Jaccard index was calculated on matching intersections of individual segmented cell. This is shown below, where *R* represents a set of pixels of a‘reference” (ground truth) cell, and *S* represents a set of pixels in a “segmented” (model predicted) cell:

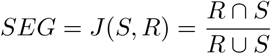

The Detection Accuracy Measure (DET) ^18^, quantifies a model’s ability to identify cells in an image without regard to the exact boundary the model produces. This is done with a graph-based approach: The ground truth and model prediction each generate their own graph, with each cell serving as a node. Then, the number of basic graph operations (e.g. add node, remove node, split node into two) required to convert the predicted graph into the ground truth graph is computed. This is referred to as the “Acyclic Oriented Graph Matching Measure for Detection” or AOGM-D.

For normalization, the number of graph operations needed to convert an empty graph into the ground truth graph is also computed. This is denoted AOGM-D_0_. Finally, a normalized metric between 0 and 1 can be computed as follows:

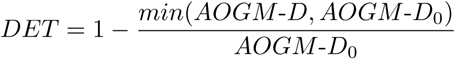

For both SEG and DET, the overall performance is defined on an interval between 0 and 1 where better performance is indicated by values closer to 1.

### 2.3 The Default Segment Anything Model

As mentioned, Segment Anything Model is a state of the art generalist model for image segmentation. A diagram of SAM’s architecture is displayed in Figure 2. SAM consists of a large image encoder, based on ViT ^15^, that converts a 1024×1024 image into a condensed embedding vector. Optionally, an input mask can be added to this embedding, as an additional input to the model. The image embedding is then supplied to a lightweight mask decoder along with an encoded prompt (or a default prompt embedding if no prompt is supplied). Subsequently, this mask decoder generates three sets of 256×256 binary masks, each accompanied by a “score” value denoting the model’s confidence for each mask. Finally, these masks are upscaled bilinearly to match the input dimension. Multiple masks are supplied to resolve ambiguity if there are multiple reasonable segmentations for a certain prompt. Meta offers pretrained variants of Segment Anything in 3 sizes, corresponding to the size of the image encoder: Base, Large, and Huge. This creates three SAM models of varying sizes based on image encoders, referred to as SAM-base, SAM-large, and SAM-huge.

**Fig. 2:**
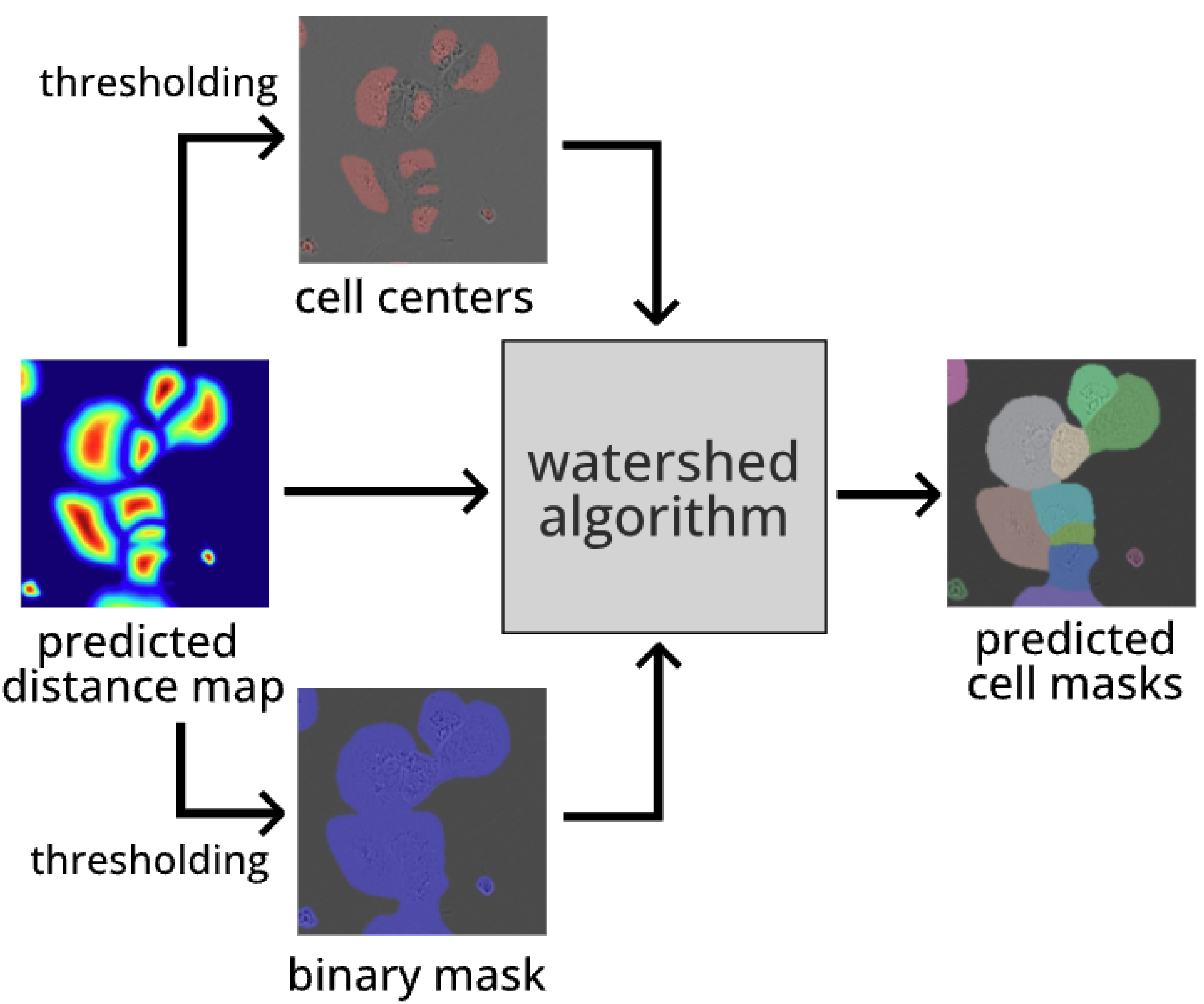
Architecture of Segment Anything Model.

Unfortunately, when employing the existing fine-tuning approaches seen in MedSAM^9^ or SAMed ^10^, difficulties arise with issues specific to the domain of cell segmentation. Using standard 2 (cell, background) or 3 (cell, cell border, background) category segmentation with the Segment Anything architecture, we observe poor results when cells have low contrast boundaries as is seen in Figure 3a. We find performance to be poor even when employing pixel-wise weighted loss functions ^6, 7^ which have been used to tackle this problem in the past. When these common characteristics arise, adjacent cells are predicted as being merged together along weak edges. Additionally, these subpar segmentation results have a detrimental impact on the accuracy of insights derived from segmentation. For instance, they can lead to an underestimation of the cell count and an inflation of the average cell size when cells are merged.

**Fig. 3:**
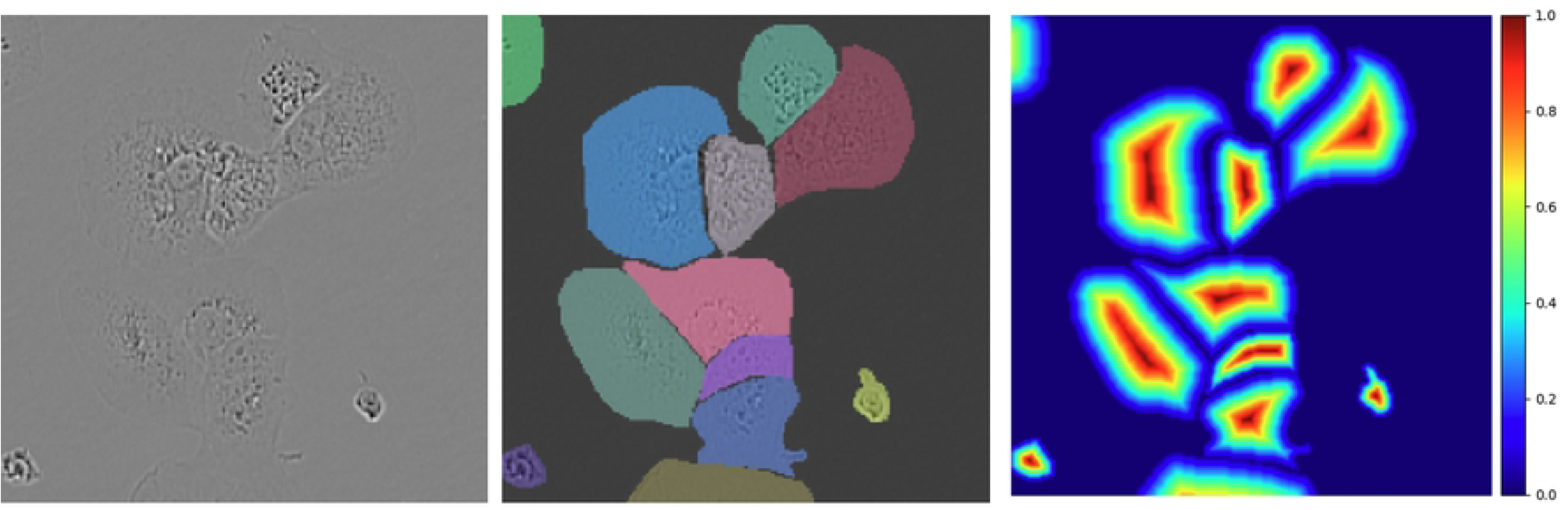
(a) A cropped microscope image from the LIVECell test set, (b) overlayed ground-truth cell masks, and (c) a distance map computed from these cell masks (right).

### 2.4 Segmentation as Regression

We formulate cell segmentation as a regression problem, drawing inspiration from Cell-Pose ^8^, to address difficulties with soft edges. Rather than predicting segmentation masks directly like U-Net ^6^, MedSAM ^9^, or SAMed ^10^, we predict a real-valued distance map, describing the euclidean distance from each pixel to its cell’s boundary (or 0 if the pixel is not part of a cell). We normalize the map such that each pixel is a real number between 0 and 1 by dividing the distance map by the max distance within each cell. An example of an annotation and its associated distance map is shown in Figures 3b and 3c respectively. Existing functions for efficiently producing these distance maps are present in common Python packages like SciPy ^19^ and OpenCV ^20^. By predicting these distance maps instead of discrete categories, this problem of merged cells can be reduced.

This approach does have some drawbacks for cells consisting of multiple compartments. Namely, when a cell is peanut-shaped (as would be the case if it was close to splitting), the distance map looks very similar to that of two disjoint cells. This makes it difficult to recover the individual cell boundaries and the cell may be incorrectly measured as two separate cells. For other non-circular shapes (like ovoid or “C-shaped” cells) we observe through empirical testing that this postprocessing technique can properly recover these geometries.

We compute a distance map using each annotation mask in the training set before training time. Because these images are invariant or almost invariant to all our data augmentation strategies, we can avoid recomputing these maps in every training step.

### 2.5 Architecture

An architecture diagram for SAMCell is shown in Figure 4. Because of VRAM constraints, concerns with inference time, and SAM’s modest performance improvements with larger models, we choose to fine-tune SAM-base.

**Fig. 4:**
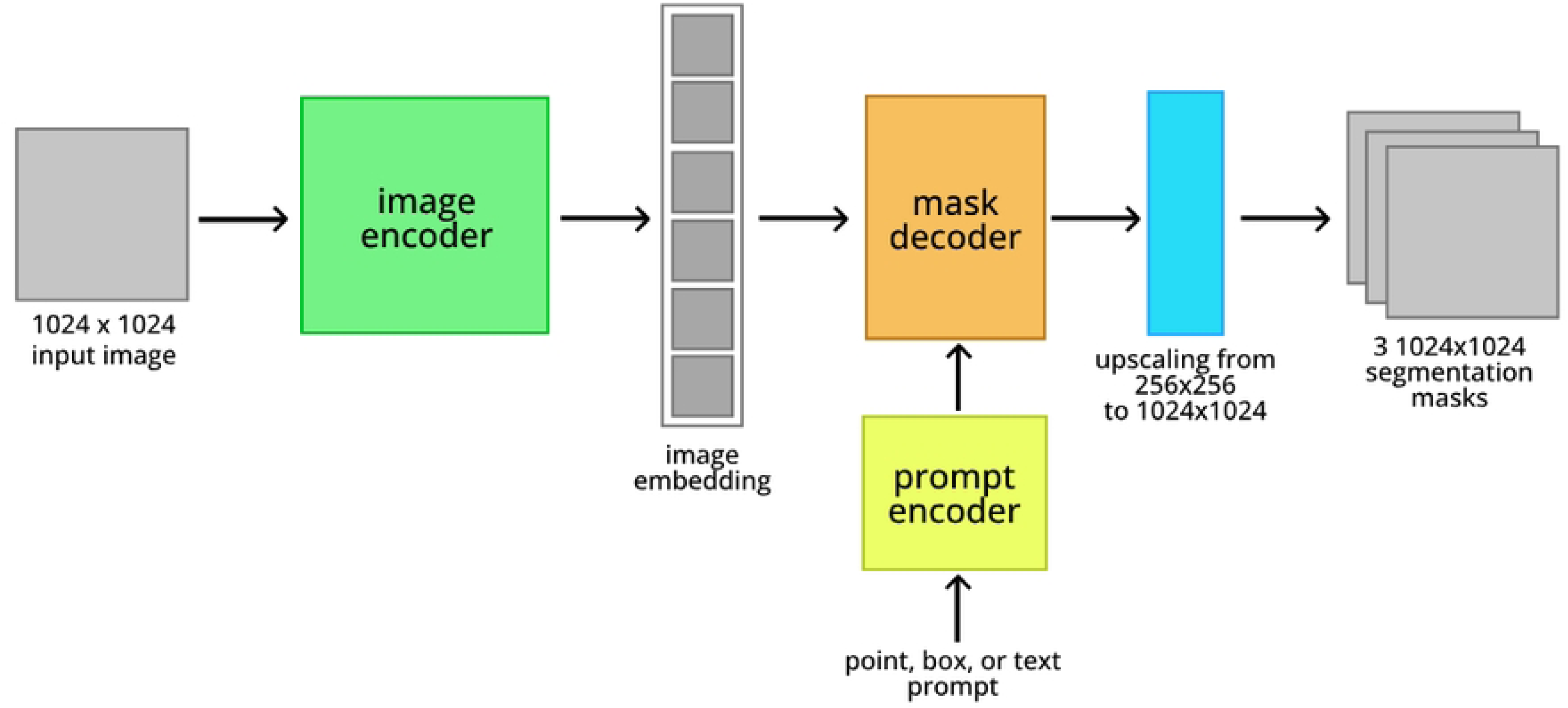
Architecture of SAMCell.

#### Pre-Processing

We find that applying Contrast Limited Adaptive Histogram Equalization (CLAHE) ^21^ is a beneficial preprocessing step. This approach eliminates any nonuniformities in brightness across the microscope field, and increases the visibility of hard to distinguish edges. Subsequently, following SAM ^5^, we normalize all input images to zero-mean and unit-variance.

#### Sliding Window Approach

To inference SAMCell, we first separate an input image into a series of 256×256 patches using a sliding window approach, like in U-Net ^6^. We use an overlap of 32 pixels on either side of these patches, to avoid poor classification of areas partially in frame. To convert each 256×256 patch to the 1024×1024 input size of the image encoder, we use bilinear upsampling. Then, we pass each image through our fine-tuned SAM, producing a distance map prediction. We apply the sigmoid activation function to the mask decoder output such that the output (predicted distance map) is in the correct range: [0, 1].

#### Eliminating Prompt

Manually supplied prompts for densely packed cells are difficult and laborious to produce, as cells in culture often number in the hundreds per image. As such we forgo SAM’s default prompting capability. To accomplish this, we employ an approach similar to that used by SAMed ^10^. We freeze the prompt encoder during fine-tuning and always input SAM’s default prompt. Because the prompt embedding is static over fine-tuning, the mask decoder learns to predict the distance map from the image embedding exclusively.

#### Finetuning Methodology

Drawing inspiration from previous SAM fine-tuning approaches ^9, 10^, we fine-tune all parameters in SAM’s lightweight mask decoder. We deviate from prior work ^10^ by finetuning all parameters in the image encoder. Because we aim to inference without user-supplied prompts, we freeze the prompt encoder.

#### Post-Processing

Post-processing must occur to convert a predicted distance map into predicted cell masks. An outline of this process is shown in Figure 5. First, via thresholding the predicted distance map, we create two binary images. The first image, we denote “binary mask”, tells whether a certain pixel belongs to a cell or background. Empirically, we find that distance map *>* 0.05 yields a good binary mask. The second image, we denote “cell centers”, must contain exactly one connected component per cell, someplace in the body of the cell. We find that distance map *>* 0.5 produces good masks for this. Fortunately, we find that these threshold values are not sensitive to the image being inferenced or the training set used. As such, we use these same threshold values for all experiments.

**Fig. 5:**
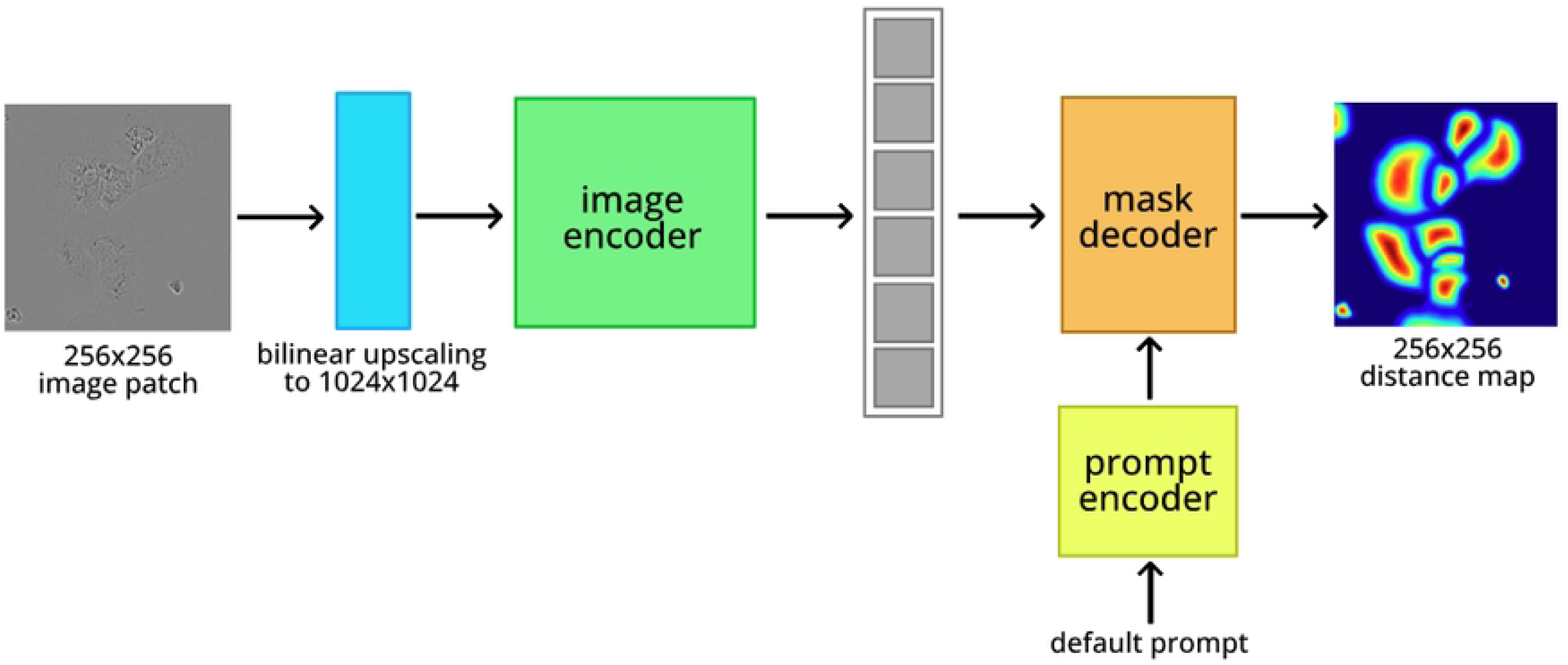
Post-processing procedure used to recover a set of distinct cell masks from a SAMCell distance map prediction.

We apply the Watershed algorithm ^22^, a classical approach to boundary detection, to recover cell masks from the distance map. The idea of this algorithm is inspired by how water floods a 3D region: first collecting at the lowest values in an image, then rising to higher regions. As the water rises higher, previously separate pools of water join together along a boundary line - the predicted boundary of the cell. As with producing the distance map, existing high-speed implementations of this algorithm exist in common libraries like OpenCV ^20^ and scikit-learn ^23^.

### 2.6 Data Augmentation

To encourage generalization, we employ data augmentation during training as follows:

- random horizontal mirroring
- random rotation between −180° and 180°
- random rescaling between 80% and 120% of the original image dimensions (preserving aspect ratio)
- random brightness adjustment between 95% and 105% of original
- randomly invert the image

Although not often seen in prior works, we find random inversion of the image to help encourage generalization. Certain microscopy approaches (e.g. dark field) show bright cells on a darker background. Other approaches (e.g. phase contrast) yield images with cells as a darker color on a brighter background. We use random inversion to create invariance towards this difference, with the aim of creating a model that can interpret images from a range of microscopy methods.

Once data augmentation has been applied to an image, a random 256×256 patch of the image and associated distance map (augmented by mirroring, rotation and rescaling only) is used for fine-tuning. As such, every epoch, SAMCell is trained on one unique, augmented 256x256 patch from every image.

### 2.7 Training Protocol

We train SAMCell for 40 epochs on an Nvidia RTX 4090 GPU, requiring about 8 hours of training time. We start with the pretrained SAM-base model and employ the AdamW ^24^ optimizer with an initial learning rate of 0.0001, and a weight decay of 0.1. We set *β*_1_ to 0.9 and *β*_2_ to 0.999. Following SAMed ^10^, we employ a learning rate warm-up ^25, 26^ with a period of 250 and a linear decrease in learning rate to 0 over the training period. Due to GPU VRAM limitations, we use a batch size of 2 for training. Because we pose the task as a regression to a distance map, we train with L2 loss.

## 3 Results and Discussion

### 3.1 Datasets

We aim to evaluate both test-set and zero-shot performance of our model. As shown in Figure 6, the zero-shot datasets we create differ in cell type, microscope contrast, and field brightness, compared to a sample image from the Cytoplasm training dataset. Thus, we argue these datasets act as a realistic evaluation of model generalization.

**Fig. 6:**
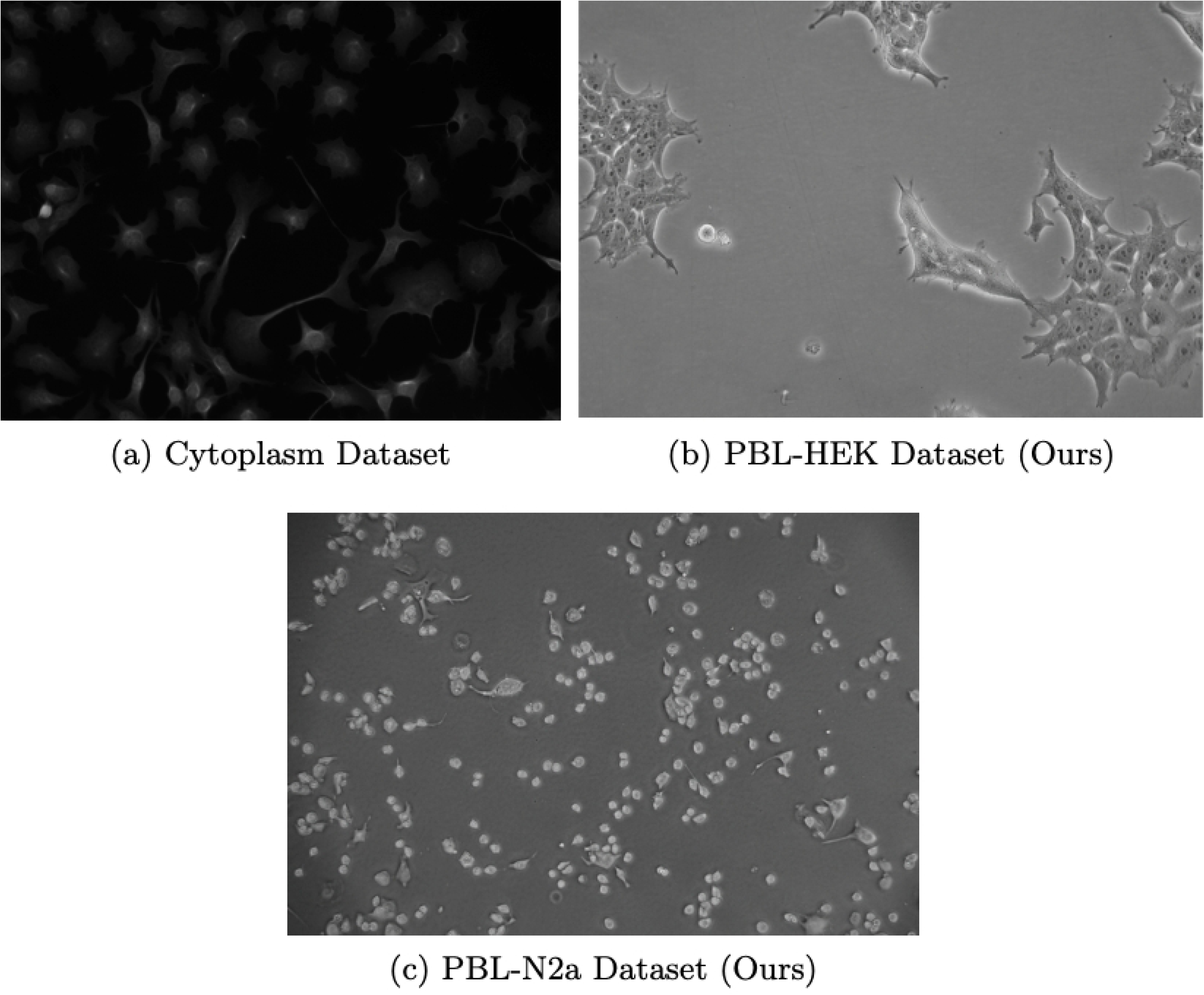
(a) Example image from the Cytoplasm dataset used for training and example images from each of our two zero-shot datasets, (b) PBL-HEK and (c) PBL-N2a.

### 3.2 Comparison to Default SAM

Although the off-the-shelf Segment Anything Model demonstrates impressive performance on a variety of tasks, we observe some common failure modes that can limit its usefulness to microscopy. We illustrate these downfalls in Figure 7. When attempting to segment densely packed cells in “automatic mask generation” mode (sampling prompt points in a uniform grid), SAM can often segment large clumps, rather than individual cells. Further, we find that SAM is prone to segmenting empty portions of the cover slip. Although this shortcoming can be overcome by a biologist prompting SAM for every cell, this introduces a manual, tedious process.

**Fig. 7:**
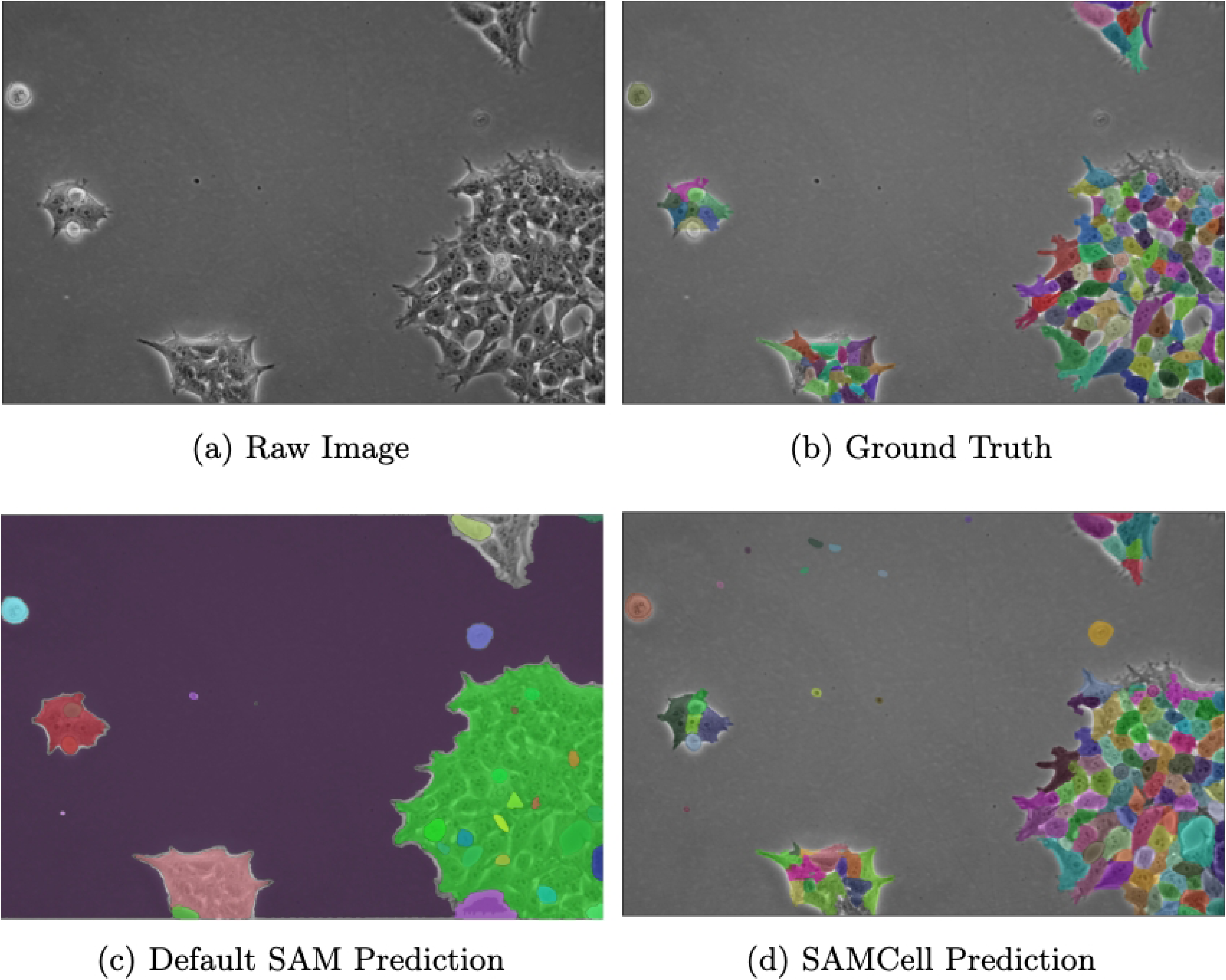
(a) A single Phase Contrast image of HEK cells, (b) annotated manually by an expert, (c) automatically annotated by Meta’s SAM-huge model, and (d) automatically annotated by our SAMCell model. The above image is from our PBL-HEK zero-shot dataset.

These failures likely stem from SAM’s training set, which, although large in scale, is focused on everyday images like segmenting food in a supermarket or people in a sporting event ^5^. Cell segmentation is a unique task: cells routinely have irregular shapes and are often clumped together with difficult to discern boundaries. As such, SAM does not have a strong prior for the appearance of cells because of its more generic dataset. This leads to SAM predicting boundaries in images that often do not correspond to individual cells.

Our fine-tuning of Segment Anything effectively resolves this shortcoming. During fine-tuning, the model was trained on a large corpus of cell masks in microscopy images. This improves its performance on related tasks, like the image in Figure 7d.

### 3.3 Baseline Methods

In addition to the default Segment Anything Model, we select 3 existing cell segmentation models as baselines for comparison:

*Stardist* ^27^: Stardist predicts cell shape as series of line segments radiating from the cell’s center point. These line segments are evenly spaced around the center and variable in length. The ends of these line segments are connected to form the boundary of a cell. This approach leverages a modified U-Net architecture to predict the component line segments for segmentation. We train this model for 50 epochs on each dataset, with the default 32 line segments per cell.

*Cellpose* ^8^: Cellpose attempts to predict specialized x and y gradients from an input image. Then, these gradients are combined to produce a smooth increase in value from cell center to the edges. The watershed algorithm is used to recover cell boundaries from these gradients. Like Stardist, a modified U-Net architecture is used as a backbone for predictions of, in this case, image gradients. Pretrained models from the Cellpose authors exist for both the LIVECell and Cytoplasm datasets. As such, we simply use these weights without modification^a^.

*CALT-US* ^28^: To provide an additional point for comparison, we select a high performing model from the Cell Tracking Challenge ^17^. This model uses a novel loss function, adding a regularization term to cross entropy loss with the goal of improving model predictions around the boundaries of cells. Using this loss function, the authors train a U-Net ^6^ to classify an each pixel in image into one of four categories: background, cell, or cells touching and gap between cells. Finally, the authors post-process the U-Net output to obtain cell boundaries. Conveniently, the authors provide an open-source implementation in a software package called jcell^b^. For the baseline used in this paper, we train for 100 epochs with batch size 32 and all the default parameters in jcell.

### 3.4 Test-Set Performance

As shown in Table 1, SAMCell demonstrates strong test-set performance, surpassing baseline performance on both the LIVECell and Cellpose Cytoplasm test sets. We hypothesize this success can be attributed to two key factors: Firstly, SAMCell inherits SAM’s image encoder which is pretrained on a diverse dataset of 11 million diverse, everyday-life images. This pretraining provides SAMCell with a strong prior for objects in general, helping the model to better detect features that contribute to a cell boundary while ignoring extraneous features like field brightness or microscope contrast.

**Table 1:** A performance comparison between our fine-tuned model, SAMCell and 3 existing cell segmentation baselines.

|  | SAMCell (ours) | Stardist | Cellpose | CALT-US |
| --- | --- | --- | --- | --- |
| Dataset: *Livcell* |  |  |  |  |
| SEG | 0.651876 | 0.572233 | 0.588850 | 0.559680 |
| DET | 0.892543 | 0.770644 | 0.778827 | 0.789718 |
| Dataset: *Cellpose Cytoplasm* |  |  |  |  |
| SEG | 0.611144 | 0.557178 | 0.579542 | 0.415172 |
| DET | 0.865908 | 0.774016 | 0.748702 | 0.571518 |

Secondly, SAMCell boasts a larger parameter count compared to our U-Net-backed baselines. This allows our model to better fit large training sets, like LIVECell and

Cytoplasm, without underfitting. Further, we speculate this higher parameter count allows the model to better capture complex features, like low-contrast cell borders or tightly packed clumps of cells.

### 3.5 Zero-Shot Performance

As with test-set evaluations, we notice impressive zero-shot capabilities with SAMCell seen in Table 2. We observe our method besting baseline approaches, for both our PBL-HEK and PBL-N2a datasets. We speculate that much of this advantage comes from SAM’s significant pretraining corpus. SAM pretraining provides SAMCell with prior knowledge of boundaries in non-microscopy images, which may be helpful in determining cell boundaries. This idea is further supported by our ablation study in Section 3.6.

**Table 2:** A zero-shot performance comparison between SAMCell trained on the LIVE-Cell dataset (SAMCell-LIVECell), SAMCell trained on the Cellpose Cytoplasm dataset (SAMCell-Cyto) and baselines trained on the Cellpose Cytoplasm dataset.

|  | SAMCell-Cyto | SAMCell-LIVECell | Stardist | Cellpose | CALT-US |
| --- | --- | --- | --- | --- | --- |
| Dataset: *PBL-HEK* |  |  |  |  |  |
| SEG | **0.364487** | 0.356793 | 0.141827 | 0.252739 | 0.070721 |
| DET | **0.728183** | 0.707649 | 0.235729 | 0.387731 | 0.161191 |
| Dataset: *PBL-N2a* |  |  |  |  |  |
| SEG | **0.701262** | 0.405612 | 0.596675 | 0.642347 | 0.458429 |
| DET | **0.936989** | 0.627771 | 0.851225 | 0.885298 | 0.632555 |

Interestingly, SAMCell trained on the Cellpose Cytoplasm dataset outperforms SAMCell trained on the LIVECell dataset on both of our zero-shot datasets. This is despite the LIVECell dataset containing a much larger samples size (∼ 500 vs ∼ 3000 images in each respective training set). This could be due to a higher diversity of microscopy images in the Cytoplasm Dataset as it was produced via scraping the internet. This results in images which come from a verity of cell lines and imaging techniques, compared to only eight cell lines imaged via phase contrast in LIVECell. Because of the stronger zero-shot performance of the SAMCell variant trained on the Cytoplasm dataset, we opt to show zero-shot baseline methods with cytoplasm pretraining in Table 2.

Across all approaches, we observe much higher values for PBL-N2a compared to PBL-HEK. We attribute this to differences in appearance between the two cell lines. Neuro-2a (N2a) cells contain a circular morphology with high contrast borders, as shown in Figure 6c. Further, these cells are less likely to grow in tightly packed clumps. Conversely, Human Embryonic Kidney (HEK) 293 cells are prone to grow in more densely packed regions and are less circular in form, as seen in 6b. These differences result in PBL-HEK producing a more difficult segmentation task, and thus lower SEG and DET metrics.

### 3.6 Pretraining Ablation Study

Approaches based on the Segment Anything Model naturally inherit pretrained weights from the extensive dataset used to train SAM. To gauge pretraining’s impact, we conduct an ablation study. We compare SAMCell as described, finetuned starting from Meta’s pretrained weights ^5^, to a SAMCell variant trained starting from random weights. Test set performance on both datasets is shown in Table 3. Note the significant drop in performance when random weight initialization is used instead of pretrained weights, suggesting pretraining has a strong impact on SAMCell’s performance.

**Table 3:** Performance of SAMCell starting from pretrained weights (SAMCell) compared to starting from a random weight initialization (SAMCell-rand).

|  | SAMCell | SAMCell-rand |
| --- | --- | --- |
| Dataset: *Livcell* |  |  |
| SEG | **0.651876** | 0.106720 |
| DET | **0.892543** | 0.510541 |
| Dataset: *Cellpose Cytoplasm* |  |  |
| SEG | **0.611144** | 0.132846 |
| DET | **0.865908** | 0.430565 |

### 3.7 Limitations

The main strength of the Segment Anything Model architecture is its large, ViT based image encoder, but this increases computational requirements compared to other lighter weight baselines. Because of the model’s size, it can capture complex features and achieve high performance on large and diverse datasets - in both the original training set ^5^, and the microscopy datasets used in this work. Unfortunately, running inference on a larger model is more computationally expensive. With a high-end consumer GPU, the Nvidia RTX 4090, we find that running SAMCell on a single image takes about 5 seconds. We find that under the same conditions, our baselines inference in well under a second. This is likely because our baselines are backed by the much slimmer U-Net architecture which requires reduced computational resources to run.

We find this difference is exacerbated when a GPU is not available as would be the case on lower-end computers. We find that running SAMCell on CPU (an AMD Ryzen 7 7700X) takes approximately 2 minutes and 20 seconds per image. We observe that our lighter-weight baselines are able to inference in a just few seconds under these conditions. As such, we recommend running SAMCell on a higher-end computer with a GPU to mitigate this limitation.

### 3.8 User Interface

To make SAMCell more accessible to users without a machine learning background, we create a user-friendly front-end for SAMCell. Using this, users can click and drag microscopy images into a straightforward window and see results from SAMCell’s automatic segmentation. The workflow of the user interface is shown in Figure 8. The landing page invites a user to drag and drop microscopy images as shown in Figure 8a. Then, as shown in Figure 8b, users can select and view images to process with SAMCell. Images in the list that have been processed are denoted green, and images being currently processed are highlighted in yellow. After an image is processed, segmentation masks can be visualized as seen in Figure 8c. Finally, we supply a few relevant metrics extracted from the segmentation result to aid biologists in culturing cells: number of cells detected, average cell area, confluency, and number of neighbors for each cell. A table with this information is presented to the user, seen in Figure 8d. As mentioned in Section 3.7, we recommend a computer with a GPU to use SAMCell, to ensure images are processed quickly. If a local GPU is not available, we recommend using a cloud GPU via our Google Colab notebook^c^, though this takes slightly more expertise than a simple GUI.

**Fig. 8:**
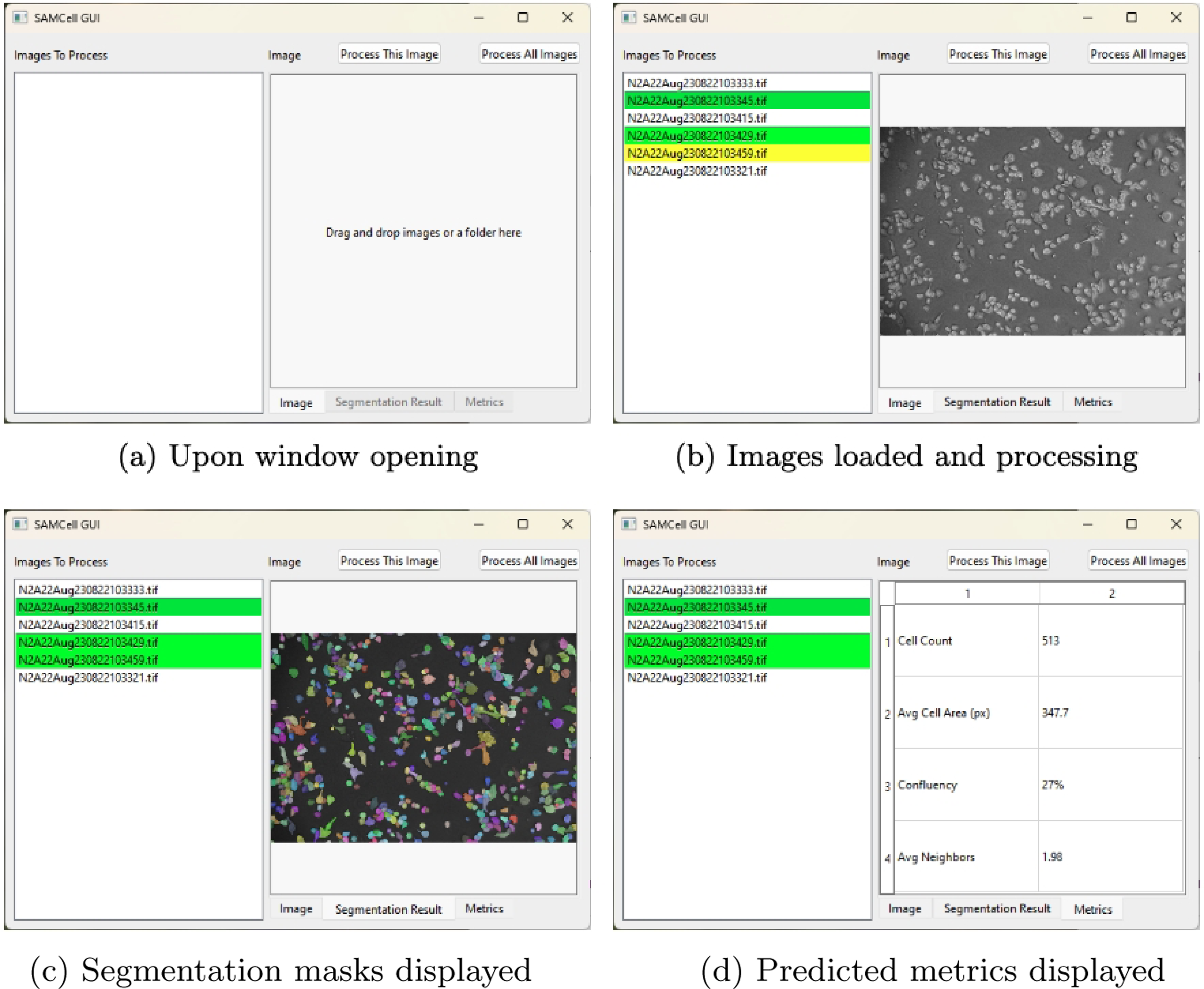
A demonstration of SAMCell’s interface throughout the various stages of use.

## 4 Conclusion

We present SAMCell, a cell segmentation model based on the Segment Anything Model architecture. Using our model, we observe state of the art performance exceeding powerful and commonly cited baselines like Cellpose and Stardist. We evaluate our model for two main use cases: when a training dataset for a similar task is available (test-set performance) and when no training dataset is available for the specific task (zero-shot performance). In both of these circumstances, our approach exceeds existing models in quantitative evaluation. To make SAMCell more accessible, we present a user-friendly interface to make this powerful model available to biological researchers without a machine learning background. We leave for future work and further exploration of our post-processing methodology. Potential projects include applying our approach to a lighter-weight model like a U-Net and comparing performance to SAMCell.

We supply training and evaluation code^d^ for SAMCell, as well as our user interface^e^ on Github. Additionally, we provide the trained weights for SAMCell, as well as our custom datasets (PBL-HEK and PBL-N2a).^f^

## 5 Declarations

### 5.1 Ethics approval and consent to participate

Not applicable.

### 5.2 Consent for publication

Not applicable.

### 5.3 Competing interests

None to declare.

### 5.4 Sourcing of Cells

The cells imaged in this paper were originally purchased from Sigma Aldrich in 2021. Subsequently, the vials used for this work were brought back up from deep freeze in May of 2023.

### 5.5 Availability of data and materials

The LIVECell ^16^ and Cellpose Cytoplasm ^8^ datasets we use for training are available from their respective authors. Code for training and testing SAMCell is available on GitHub. ^g^ Our zero-shot datasets (PBL-N2a and PBL-HEK), as well as trained model weights, are available on the same repo.^h^ Code for our GUI is hosted also hosted on GitHub at a different repository.^i^

### 5.6 Funding

The authors acknowledge the NIH BRAIN Initiative Grant (NEI and NIMH 1-U01-MH106027-01), NIH R01NS102727, NIH Single Cell Grant 1 R01 EY023173, NIH R01DA029639 and NIH RF1AG079269, support from Georgia Tech through the Institute for Bioengineering and Biosciences, Invention Studio, and the George W. Woodruff School of Mechanical Engineering.

## 6 Availability and Requirements

**Project name:** SAMCell

**Project home page:**

Model Training and Evaluation: https://github.com/NathanMalta/SAMCell/ User Interface: https://github.com/NathanMalta/SAMCell-GUI

**Operating system(s):** Development was conducted on Ubuntu 22.04. The software can run on any operating system supporting Python, including Windows, Mac, and Linux.

**Programming language:** Python 3.11

**Other requirements:** Hardware: A computer with a powerful GPU is recommended for optimal performance. Software: Common deep learning packages like PyTorch and NumPy are used by this project. A full list of required libraries is present in each repository’s requirements.txt.

**License:** MIT License

**Restrictions to use by non-academics:** None

## Footnotes

a Cellpose pretrained models are available at https://cellpose.readthedocs.io/en/latest/models.html

b https://jcell.org/

c https://colab.research.google.com/drive/1016jr1JTtSI4kUIaHmnmXn290n-SM2eP?usp=sharing

d https://github.com/NathanMalta/SAMCell

e https://github.com/NathanMalta/SAMCell-GUI

f https://github.com/NathanMalta/SAMCell/releases/tag/v1

g https://github.com/NathanMalta/SAMCell

h https://github.com/NathanMalta/SAMCell/releases/tag/v1

i https://github.com/NathanMalta/SAMCell-GUI

## Notes

### Competing Interest Statement

The authors have declared no competing interest.

## References

1. Verma A, Verma M, Singh A. Chapter 14 - Animal tissue culture principles and applications. In: Verma AS, Singh A, editors. Animal Biotechnology (Second Edition). second edition ed. Boston: Academic Press; 2020. p. 269-93. Available from: https://www.sciencedirect.com/science/article/pii/B9780128117101000124.

2. Cell Culture Basics Handbook. Waltham, MA: Thermo Fisher Scientific; 2020.

3. Jindal D, Singh M. In: Counting of Cells. Cham: Animal Cell Culture: Principles and Practice, pp. 131–145. Springer International Publishing; 2023. Available from: 10.1007/978-3-031-19485-69.

4. Weigert M, Schmidt U. Nuclei Instance Segmentation and Classification in Histopathology Images with Stardist. In: 2022 IEEE International Symposium on Biomedical Imaging Challenges (ISBIC); 2022. p. 1-4.

5. Kirillov A, Mintun E, Ravi N, Mao H, Rolland C, Gustafson L, et al. Segment Anything. 2023 IEEE/CVF International Conference on Computer Vision (ICCV). 2023:3992-4003. Available from: https://api.semanticscholar.org/CorpusID:257952310.

6. Ronneberger O, Fischer P, Brox T. U-Net: Convolutional Networks for Biomedical Image Segmentation. In: Navab N, Hornegger J, Wells WM, Frangi AF, editors. Medical Image Computing and Computer-Assisted Intervention MICCAI 2015. Cham: Springer International Publishing; 2015. p. 234-41.

7. Guerrero-Pena FA, Fernandez PDM, Ren TI, Yui M, Rothenberg E, Cunha A. Multiclass Weighted Loss for Instance Segmentation of Cluttered Cells. In: 2018 25th IEEE International Conference on Image Processing (ICIP). IEEE; 2018. Available from: 10.1109/icip.2018.8451187.

8. Stringer C, Wang T, Michaelos M, Pachitariu M. Cellpose: a generalist algorithm for cellular segmentation. Nature Methods. 2021 Jan;18(1):100–6. Available from: 10.1038/s41592-020-01018-x.

9. Ma J, He Y, Li F, Han L, You C, Wang B. Segment Anything in Medical Images. Nat Commun 15, 654. 2023.

10. Zhang K, Liu D. Customized Segment Anything Model for Medical Image Segmentation. ArXiv. 2023;abs/2304.13785. Available from: https://api.semanticscholar.org/CorpusID:258352583.

11. Hu JE, Shen Y, Wallis P, Allen-Zhu Z, Li Y, Wang S, et al. LoRA: Low-Rank Adaptation of Large Language Models. ArXiv. 2021;abs/2106.09685. Available from: https://api.semanticscholar.org/CorpusID:235458009.

12. Israel U, Marks M, Dilip R, Li Q, Schwartz M, Pradhan E, et al. A Foundation Model for Cell Segmentation. bioRxiv : the preprint server for biology; 2023..

13. Archit A, Nair S, Khalid N, Hilt P, Rajashekar V, Freitag M, et al. Segment Anything for Microscopy. bioRxiv. 2023. Available from: https://www.biorxiv.org/content/early/2023/08/22/2023.08.21.554208.

14. Hörst F, Rempe M, Heine L, Seibold C, Keyl J, Baldini G, et al.. CellViT: Vision Transformers for Precise Cell Segmentation and Classification; 2023.

15. Kolesnikov A, Dosovitskiy A, Weissenborn D, Heigold G, Uszkoreit J, Beyer L, et al.. An Image is Worth 16x16 Words: Transformers for Image Recognition at Scale; 2021.

16. Edlund C, Jackson TR, Khalid N, Bevan NJ, Dale T, Dengel AR, et al. LIVE-Cell—A large-scale dataset for label-free live cell segmentation. Nature Methods. 2021;18:1038-1045. Available from: https://api.semanticscholar.org/CorpusID:237366120.

17. Maška M, Ulman V, Svoboda D, Matula P, Matula P, Ederra C, et al. A benchmark for comparison of cell tracking algorithms. Bioinformatics. 2014 02;30(11):1609-17. Available from: 10.1093/bioinformatics/btu080.

18. Matula P, Maška M, Sorokin DV, Matula P, Ortiz-de Soĺorzano C, Kozubek M. Cell Tracking Accuracy Measurement Based on Comparison of Acyclic Oriented Graphs. PLOS ONE. 2015 12;10(12):1-19. Available from: 10.1371/journal.pone.0144959.

19. Virtanen P, Gommers R, Oliphant TE, Haberland M, Reddy T, Cournapeau D, et al. SciPy 1.0: Fundamental Algorithms for Scientific Computing in Python. Nature Methods. 2020;17:261–72.

20. Bradski G. The OpenCV Library. Dr Dobb’s Journal of Software Tools. 2000.

21. Pizer SM, Johnston RE, Ericksen JP, Yankaskas BC, Muller KE. Contrast-limited adaptive histogram equalization: speed and effectiveness. In: [1990] Proceedings of the First Conference on Visualization in Biomedical Computing; 1990. p. 337-45.

22. Beucher S, Lantúejoul C. Use of Watersheds in Contour Detection. vol. 132; 1979..

23. Pedregosa F, Varoquaux G, Gramfort A, Michel V, Thirion B, Grisel O, et al. Scikit-learn: Machine Learning in Python. Journal of Machine Learning Research. 2011;12:2825–30.

24. Loshchilov I, Hutter F. Decoupled Weight Decay Regularization. In: International Conference on Learning Representations; 2017. Available from: https://api.semanticscholar.org/CorpusID:53592270.

25. He K, Zhang X, Ren S, Sun J. Deep Residual Learning for Image Recognition. 2016 IEEE Conference on Computer Vision and Pattern Recognition (CVPR). 2015:770-8. Available from: https://api.semanticscholar.org/CorpusID:206594692.

26. Xiong R, Yang Y, He D, Zheng K, Zheng S, Xing C, et al. On Layer Normalization in the Transformer Architecture. ArXiv. 2020;abs/2002.04745. Available from: https://api.semanticscholar.org/CorpusID:211082816.

27. Schmidt U, Weigert M, Broaddus C, Myers G. Cell Detection with Star-Convex Polygons. In: Medical Image Computing and Computer Assisted Intervention - MICCAI 2018 - 21st International Conference, Granada, Spain, September 16-20, 2018, Proceedings, Part II; 2018. p. 265-73.

28. Guerrero Peña FA, Marrero Fernandez PD, Tarr PT, Ren TI, Meyerowitz EM, Cunha A. J Regularization Improves Imbalanced Multiclass Segmentation. In: 2020 IEEE 17th International Symposium on Biomedical Imaging (ISBI); 2020. p. 1-5.

